# EcoMorph: Universal morphological trait quantification from natural language prompts for ecological research

**DOI:** 10.64898/2026.07.10.737871

**Authors:** Edward I. Amoah, Zachary Bunch, Hannah M. Thomas, Harland M. Patch, Christina M. Grozinger

## Abstract

1. Morphological traits such as floral area and body size are fundamental to ecological research, serving as inputs for studies of pollinator–plant interactions, habitat quality, and biodiversity monitoring. However, accurately measuring these traits from images remains challenging, particularly in complex field conditions where existing tools exhibit reduced accuracy and limited generalizability across taxa.
2. We present EcoMorph, a modular morphological measurement system that leverages the Segment Anything Model 3 (SAM3) to quantify traits across diverse ecological contexts. Unlike task-specific segmentation models requiring domain-specific training data, SAM3’s prompt-based architecture enables segmentation of arbitrary biological structures from natural-language prompts, using the same underlying model across flowers, insects, and other targets without retraining. From the resulting segmentations, EcoMorph extracts three classes of measurement: area, linear dimensions, and object counts.
3. We validated EcoMorph across two ecological scales. At the intermediate scale, EcoMorph-derived floral area agreed closely with manual ImageJ measurements (R^2^ = 0.935, n = 74) under simple-background conditions and (R^2^ = 0.928, n = 58) under complex-background conditions, with valid predictions for 95% of images. At the fine scale, EcoMorph-derived insect body area was strongly correlated with hand-measured intertegular distance (r = 0.810, n = 349), capturing body-size variation across species from the small *Bombus impatiens* to the large *Xylocopa virginica*. Object counts matched manual counts almost exactly for well-separated insects in an insect box (R^2^ = 0.9997, n = 12).
4. By combining prompt-based segmentation with modular measurement, EcoMorph enables high-throughput quantification of area, size, and abundance from heterogeneous image sources without taxon-specific training. This generality supports a broad range of ecological applications, including pollinator and plant trait research, biodiversity and abundance monitoring, and allometric biomass estimation.

## 1. Introduction

Body size and other ecological traits play a fundamental role in shaping ecological processes, from individual physiology to ecosystem-level dynamics (Peters, 1986; Brown et al., 2004; Chave et al., 2014; Sohlstrom et al., 2018). Ecological traits such as bee body size influence key behaviors, including foraging range and resource acquisition (Greenleaf et al., 2007). Similarly, plant traits, such as flower area, affect pollinator attraction and visitation frequency (Galen, 2000; Thompson, 2001; Delgado et al., 2023). At the landscape level, the abundance, diversity, and spatial distribution of flowering plants shape bees’ ability to locate resources, navigate efficiently, and determine their degree of specialization on particular plant taxa (Pardee et al., 2023). These traits are measured across a wide range of spatial scales, from fine-scale measurements of bee body size (millimeters) (Hagen & Dupont, 2013a), to intermediate-scale flower area (centimeters) (Amoah, White, et al., 2025), to landscape-level patterns of floral cover and distribution (meters to kilometers) (Jha & Kremen, 2013). Despite their importance, accurately and efficiently quantifying these ecological traits remains challenging, as existing methods are often time- and resource-intensive, have a limited scope, and perform poorly under complex field conditions.

Automated image analysis has emerged as a promising approach for extracting ecological data at scale, yet existing tools face persistent challenges. For insect morphometrics, hierarchical segmentation pipelines such as MAPHIS enable detailed measurement of arthropod phenotypes but require task-specific segmentation training, limiting generalization across taxa (Mráz et al., 2024). For leaf morphology measurement, Python-based workflows using OpenCV have achieved high performance, though these approaches often require costly specialized imaging equipment, limiting field applicability (Zhu et al., 2023). Deep learning (DL) methods for plant phenotyping have also achieved >97% accuracy in identifying features under controlled greenhouse conditions (Pound et al., 2017); however, performance typically degrades in field settings with variable lighting and complex backgrounds. For landscape-scale measurements, DL-based methods for drone-derived floral surveys have shown promise, but precision (0.769) and recall (0.849) metrics indicate detection limitations in visually complex environments (Sookhan et al., 2024). A common constraint across these DL approaches is the requirement for task-specific model training. Deep Learning (DL) task-specific training requires extensive labeled datasets for each new application domain (Sohan et al., 2024), precluding generalization across the morphological diversity and complex backgrounds encountered in ecological research.

Recent advances in visual foundation models offer new possibilities for ecological and morphological measurement. Unlike task-specific models requiring extensive training data, the Segment Anything Model 3 (SAM3) accepts flexible prompts, including points, boxes, or text descriptions, to segment arbitrary objects (Carion et al., 2025). This prompt-based approach aligns naturally with ecological workflows where researchers need to measure diverse structures across species and contexts without retraining models for each application.

Here we present EcoMorph, a universal morphological trait quantification system that combines SAM3’s prompt-based segmentation with reference- and distance-based scaling in a single modular pipeline. We validate EcoMorph through ecological measurements at two spatial scales: intermediate-scale (i.e., cm) floral area, tested under simple and complex background conditions; and fine-scale (i.e, mm) insect morphometrics from museum specimens, compared with hand-measured intertegular distance.

**Figure 1.**
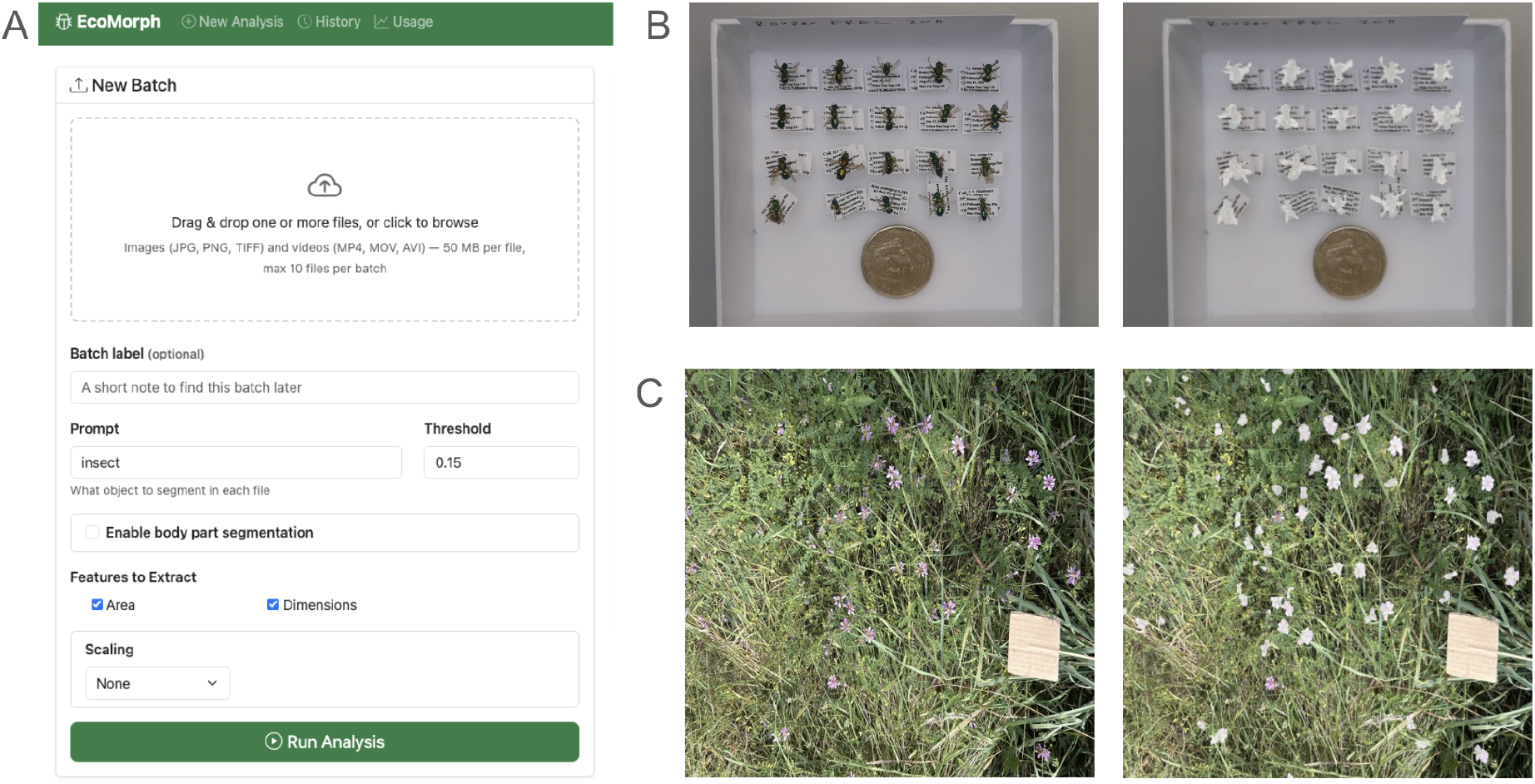
EcoMorph Application. (A) EcoMorph provides researchers with a simple, intuitive interface for measuring ecological traits from images and videos. Users can segment and measure a variety of ecological traits by adjusting the segmentation prompt and detection thresholds. Additionally, users can use the body part segmentation feature to segment pinned insects in detail. All measurements can be converted from digital pixels to real-world metrics using the scaling options. (B) EcoMorph measurement of pinned insects using a coin as the reference object for scaling. (C) Ecomorph measurement of flowers using a brown square cardboard as a reference object. You can access the software at this URL (https://mwqipf3urp.us-east-1.awsapprunner.com/)

## 2. Materials and Methods

### 2.1 System architecture overview

EcoMorph employs a modular architecture that abstracts morphological data extraction from images, enabling its application across diverse ecological systems and research contexts (Figure 2). The system comprises three primary modules arranged in a sequential pipeline: (1) the Object Segmentation Module, which identifies target structures using SAM3; (2) the Morphology Extraction Module, which computes pixel-based measurements including area, dimensions, and counts; and (3) the Measurement Scaling Module, which converts pixel measurements to physical units using either reference-based or distance-based calibration. This hierarchical design separates the processes of object detection, feature extraction, and unit conversion, allowing each component to be independently optimized or extended for specific research applications. Final outputs include area, linear dimensions, and object counts, all of which can be exported for downstream analysis, including integration with allometric equations for biomass estimation.

**Figure 2.**
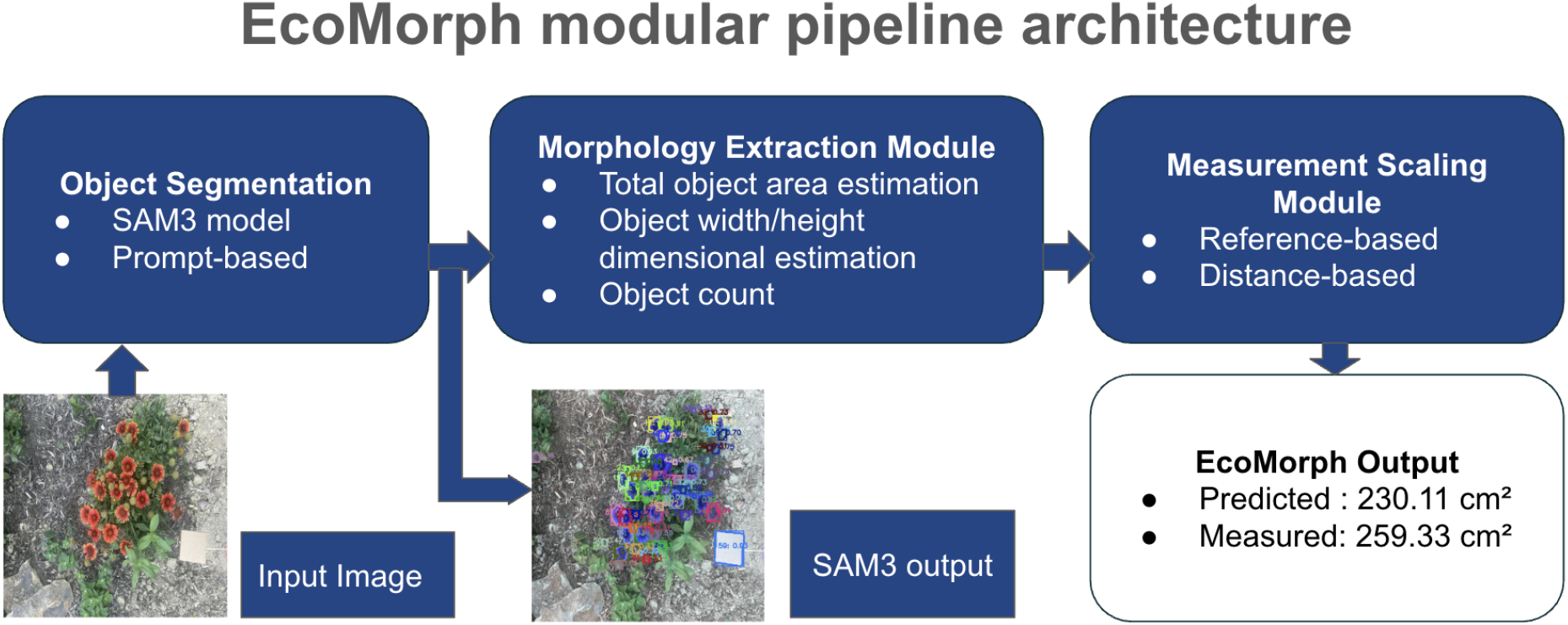
EcoMorph modular pipeline architecture for morphological trait quantification from images. The pipeline consists of three primary modules: (1) the Object Segmentation Module, which uses SAM3 for prompt-based object detection; (2) the Morphology Extraction Module, which computes pixel-based measurements including total area, linear dimensions, and object counts; and (3) the Measurement Scaling Module, which converts pixel measurements to physical units using either reference-based calibration or distance-based estimation. Final outputs include area, linear dimensions, and object counts, all of which can be exported for downstream analysis, including integration with allometric equations for biomass estimation.

#### 2.1.1 Prompt-based Object Segmentation Module

The Object Segmentation Module leverages SAM3 to identify and delineate target structures within input images. EcoMorph relies on text prompts, which can range from single nouns to short noun phrases incorporating visual descriptors such as color, shape, or size. In most cases, a single noun is sufficient: “flower,” “insect,” or “leaf” reliably segments the intended structures when the target is the dominant object in the image. When multiple candidate objects share the frame, or when the target is visually similar to surrounding elements, more specific prompts improve segmentation. For example, “pink flowers” restricts segmentation to a particular color class when multiple flowering species are present, and “brown square cardboard” disambiguates a reference object from other rectangular elements in the background. Across the experiments in this study, flowers were segmented with the prompt “flower,” insect specimens with “insect,” and reference objects with prompts matching their appearance (“brown square cardboard” for the floral experiments, “coin” for the insect experiments). This prompt-based approach eliminates the need for domain-specific training data while enabling segmentation of arbitrary biological structures, from flowers and leaves to insect bodies and tree trunks, using the same underlying model.

#### 2.1.2 Morphology Extraction Module

The Morphology Extraction Module computes measurements from segmentation masks. For area measurement, the module calculates the total pixel area by summing the areas of all pixels classified as belonging to target objects. When multiple objects are detected from a single prompt (e.g., “flower”), the module returns the aggregate area across all instances.

For dimensional metrics, the module extracts width and height measurements from segmentation masks by computing the bounding dimensions of detected objects. For single-object prompts (e.g., thorax), the module returns the width at the vertical midpoint of the mask, analogous to intertegular distance (ITD) measurements.

For count, the module counts discrete objects detected in each prompt, enabling quantification of flowers per plant, insect count per frame, and other count-based ecological metrics.

#### 2.1.3 Measurement Scaling Module

The Measurement Scaling Module converts pixel-based measurements from the Morphology Extraction Module into physical units. This module can use two alternative approaches: reference-based scaling (where a reference objective is available in the image) or distance-based scaling, which uses the distance between the camera and the object. These two approaches are described below.

##### Reference-based scaling

This approach uses a physical calibration object of known dimensions placed within the image plane. The module uses the segmented mask of the reference object, calculates its pixel dimensions, and computes a scaling factor relating pixels to physical units. This ratio is then applied to all target measurements. Reference-based scaling follows the principles of two-dimensional photogrammetry, which has been validated for estimating floral area and tree diameter, achieving high accuracy (R^2^ > 0.93) across diverse flowering plants and tree species (Amoah, McCloskey, et al., 2025; Amoah, White, et al., 2025).

##### Distance-based scaling

When physical reference objects are impractical, such as in canopy photography, drone imagery, or rapid field surveys, this sub-module calculates scaling factors from camera sensor specifications and measured subject distance from the camera. Given the sensor dimensions, focal length, and camera-to-subject distance, the module computes the physical metrics for per-pixel measurements (Luhmann et al., 2023; Sae-ngow et al., 2025).

### 2.2 Ecological validation

We evaluated EcoMorph by comparing its output against established ground-truth measurements for two ecological traits: floral area at the intermediate scale (cm) and bee body size and insect count at the fine scale (mm). Together, these validation tests whether a single prompt-based tool can deliver accurate measurements across the morphological and environmental diversity of ecological research.

#### 2.2.1 Floral area measurement validation

We validated area measurement with ImageJ, a widely used reference tool for area estimation in ecological research (Schneider et al., 2012). Validation was conducted under two conditions of progressively increasing difficulty: simple backgrounds, using top-view images of flowering plants in pots placed on a concrete floor (n=75) (data from Amoah, White, et al., 2025); and complex backgrounds, using images of flowering plants captured in outdoor garden settings under natural lighting and against mixed vegetation (n=61) (Figure 3). We fitted linear regression models to compare the EcoMorph-predicted floral area with the ImageJ-measured floral area under each of the two background conditions.

**Figure 3.**
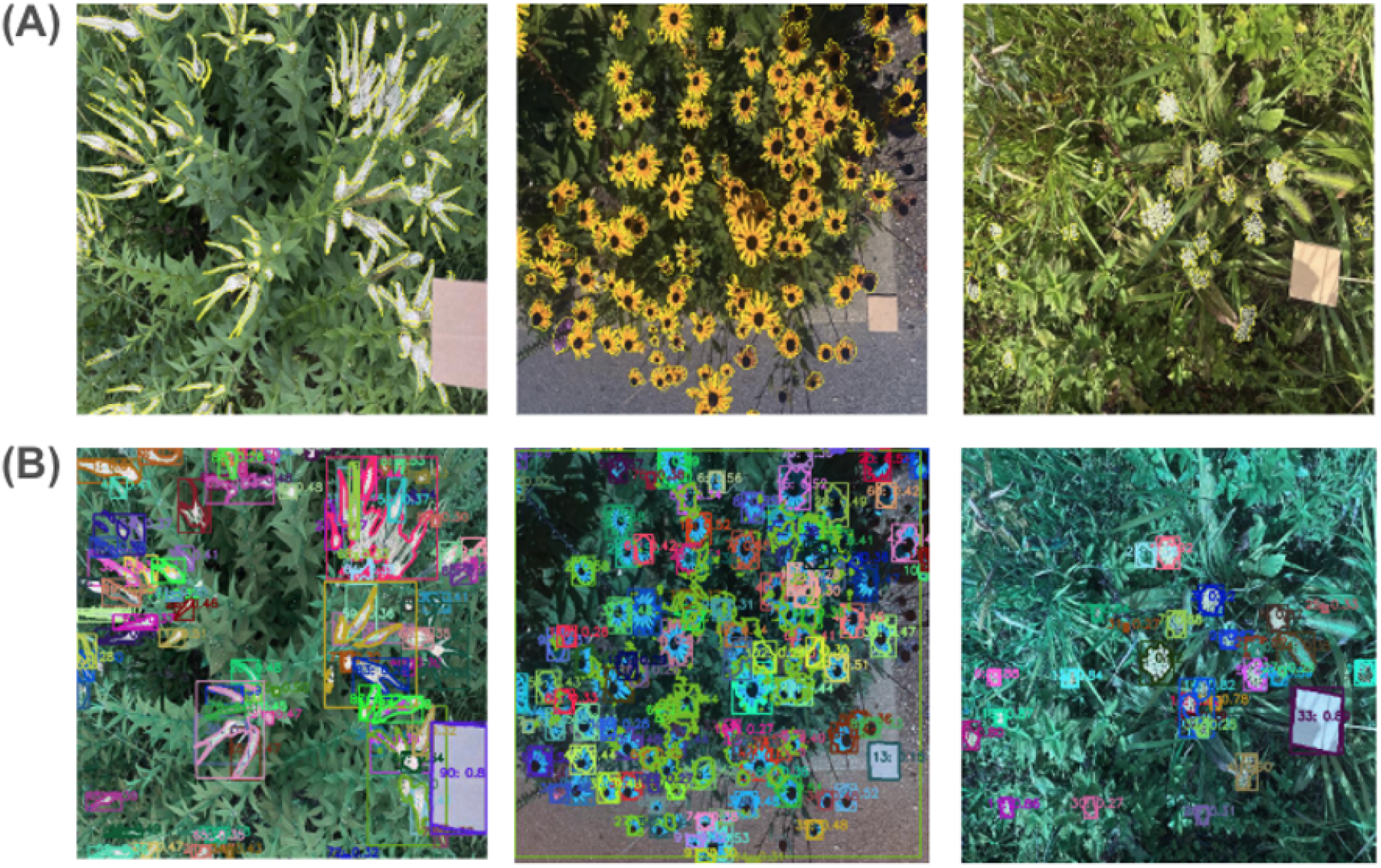
ImageJ annotations and SAM3 segmentations. (A) Manual ImageJ segmentation is shown in the top panel, and (B) SAM3 segmentation is shown in the bottom panel.

#### 2.2.2 Insect morphometric validation

To assess whether EcoMorph generalizes beyond floral measurements to fine-scale insect morphometrics, we validated its output against intertegular distance (ITD). Pinned, dry bee specimens from the Frost Entomology Museum at Pennsylvania State University were measured using an ocular micrometer mounted on a dissecting microscope to obtain ITD (Figure 4A). Specimens were then photographed using the low-cost OAK-1 (Luxonis Holding Corporation, Denver, CO, USA) camera mounted on a tripod with a United States quarter-dollar coin included in the frame as a size reference object for scale calibration (Figure 4B, C). Images were processed through EcoMorph, which automatically extracted the body area. Pearson correlation coefficients were computed between hand-measured ITD and body area for all pooled specimens spanning three species (*Bombus impatiens, Dolichovespula maculata, and Xylocopa virginica*) to evaluate performance across a broad range of body sizes.

**Figure 4.**
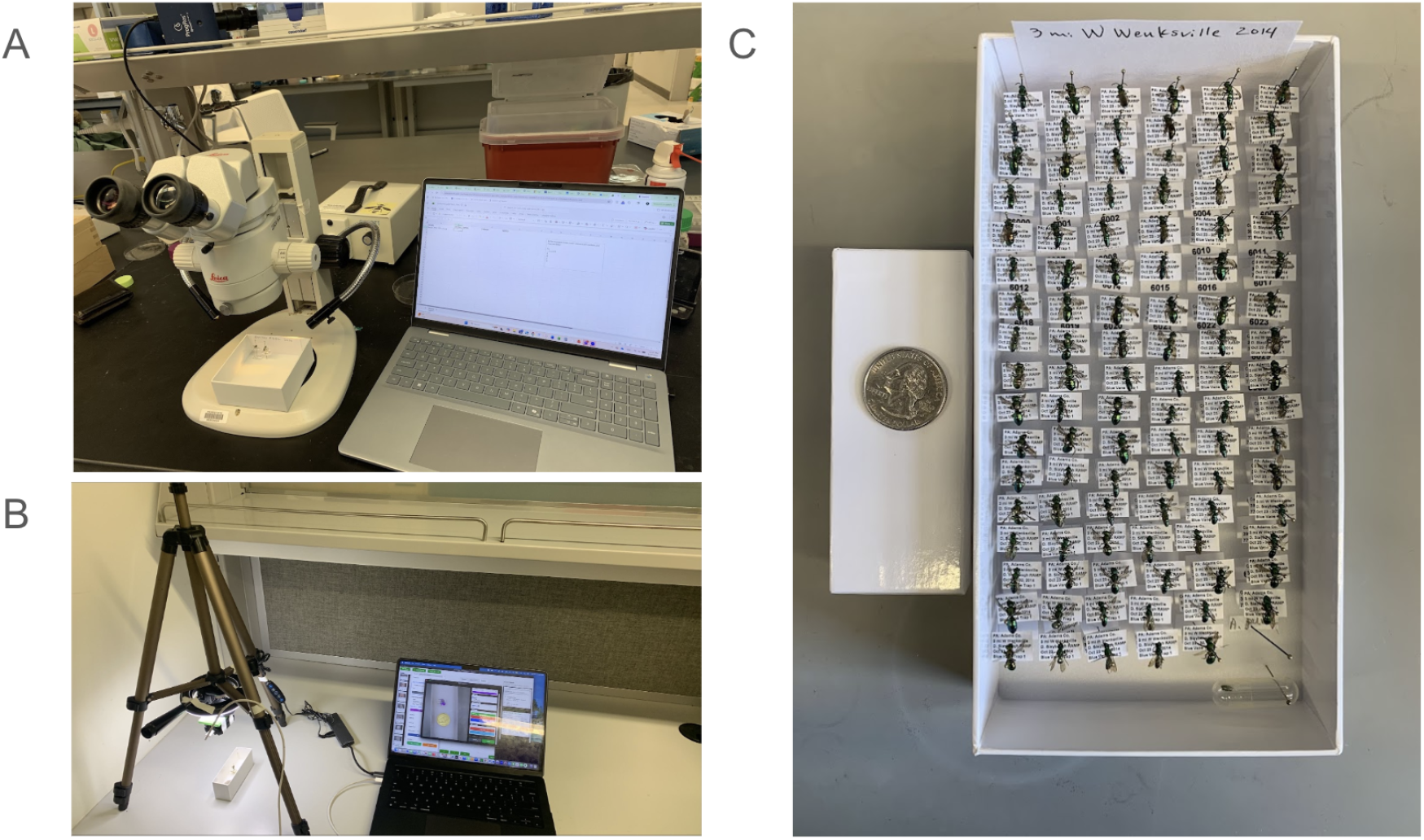
Insect morphometric validation setup. (A) Hand measurement of intertegular distance (ITD) using an ocular micrometer mounted on a dissecting microscope. (B) Photographic imaging of pinned specimens using an OAK-1 camera on a tripod, with images streamed to a laptop running EcoMorph. (C) Tray of pinned bee specimens from Frost Entomology Museum, with a United States quarter-dollar coin included in the frame as the scale reference object.

To validate EcoMorph’s count function for insects, 12 images of pinned bee specimens were photographed using the OAK-1 camera as described above. Each image contained between 7 and 80 specimens (mean 31) and was manually counted prior to processing through EcoMorph’s Morphology Extraction Module. We fitted a linear regression comparing EcoMorph-predicted counts with manual counts and quantified accuracy using root-mean-square error (RMSE) and mean absolute error.

## 3. Results

### 3.1 Floral area measurements validation

For measuring floral area on a simple background, EcoMorph estimated floral area from images with strong agreement to ImageJ ground-truth measurements (Figure 5A). A single highly influential outlier was identified using regression diagnostics and removed before analysis, resulting in a final sample size of n = 74. Predicted areas were strongly correlated with ImageJ measurements (R^2^ = 0.935, p < 0.001; Figure 5A), with a regression equation of y = −4.46 + 1.37x. Root-mean-square error was 27.79 cm^2^, with a positive bias of 19.96 cm^2^. The standard deviation of residuals was 28.17 cm^2^ (Figure 5B).

**Figure 5.**
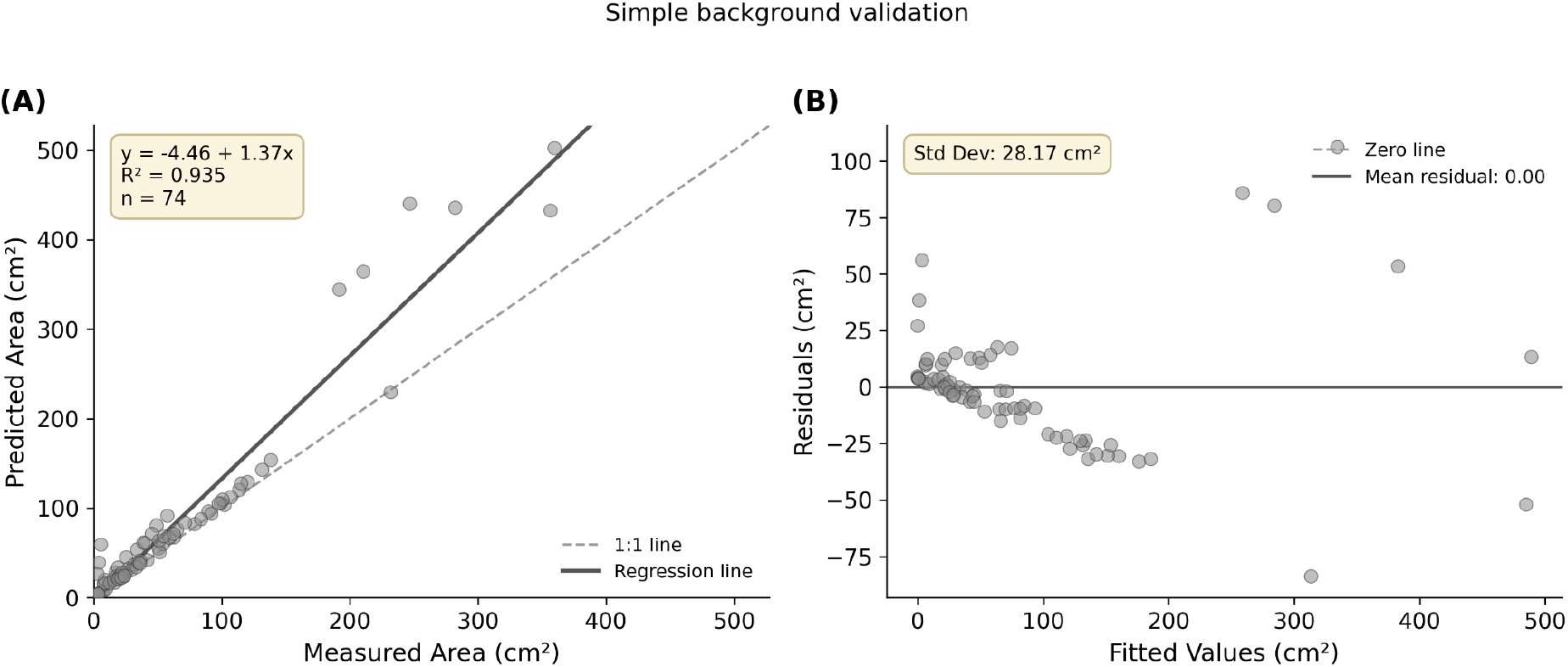
Simple background validation of EcoMorph area estimation (n = 74). (A) Relationship between EcoMorph-predicted floral area and ImageJ-measured ground truth. The solid line represents the fitted regression; the dashed line indicates the 1:1 relationship. (B) Residual plot showing prediction errors across the range of fitted values. The dashed line indicates zero residual; the standard deviation of residuals was 28.17 cm^2^.

For floral area measurement in complex field backgrounds, EcoMorph’s SAM3-based segmentation identified flowers across a range of colors and morphologies (Figure 6A). Segmentation failures occurred in three images containing fine, highly filamentous floral structures (Figure 6B). For the 58 images (i.e., 95% of the 61 evaluation images) with valid EcoMorph predictions, the system achieved R^2^ = 0.928, RMSE = 142.64 cm^2^, bias = 118.02 cm^2^, and a regression equation of y = 44.00 + 1.27x (p < 0.001; Figure 7A). The standard deviation of residuals was 145.17 cm^2^ (Figure 7B).

**Figure 6.**
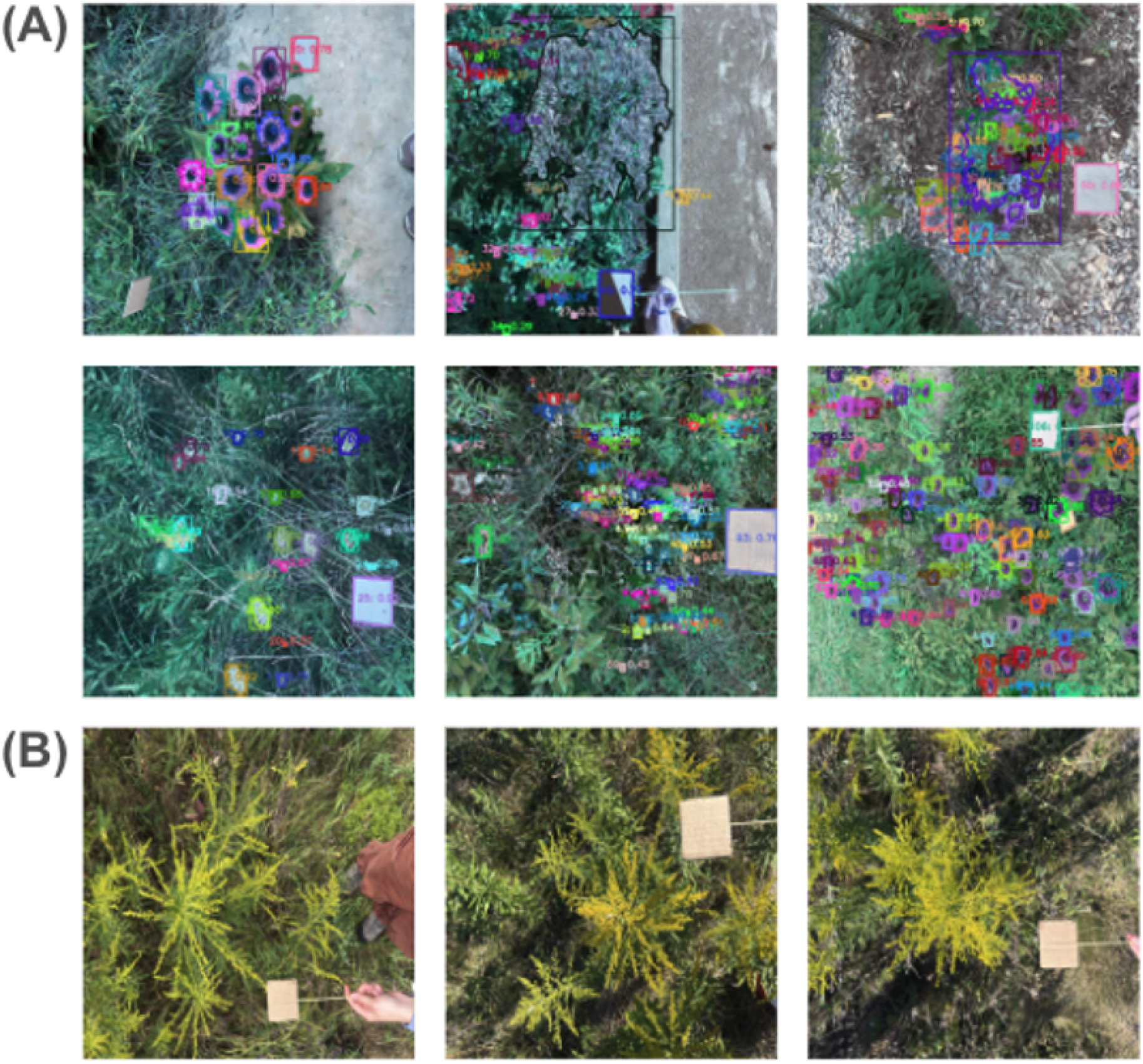
Examples of EcoMorph’s SAM3-based segmentation performance in complex background environments. (A) Successful segmentation across diverse flower colors, morphologies, and complex backgrounds. (B) Segmentation failures in images containing fine, highly filamentous floral structures.

**Figure 7.**
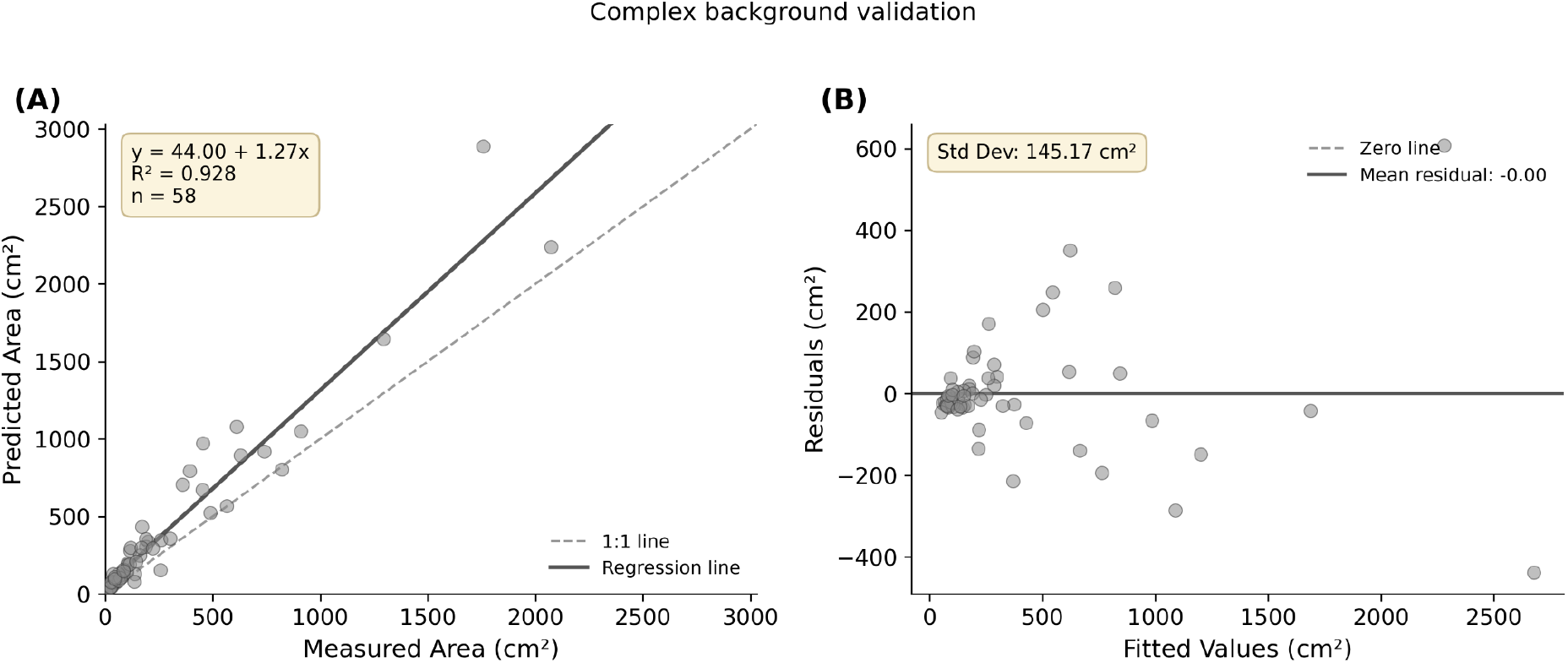
Complex background validation of EcoMorph area estimation (n = 58 of 61 images with valid predictions; 95%). (A) Relationship between EcoMorph-predicted floral area and ImageJ-measured ground truth. The solid line represents the fitted regression; the dashed line indicates the 1:1 relationship. (B) Residual plot showing prediction errors across the range of fitted values. The dashed line indicates zero residual; the standard deviation of residuals was 145.17 cm^2^.

### 3.2 Insect morphometric measurement validation

EcoMorph-derived body area was strongly correlated with hand-measured intertegular distance (ITD) across all three species (r = 0.810, n = 349; Figure 8), indicating that EcoMorph captures body size variation consistent with a widely used morphometric standard. The relationship was positive and approximately linear across the full range of sampled sizes, from small Bombus impatiens to large Xylocopa virginica, suggesting the tool performs reliably across species with generally different body sizes.

**Figure 8.**
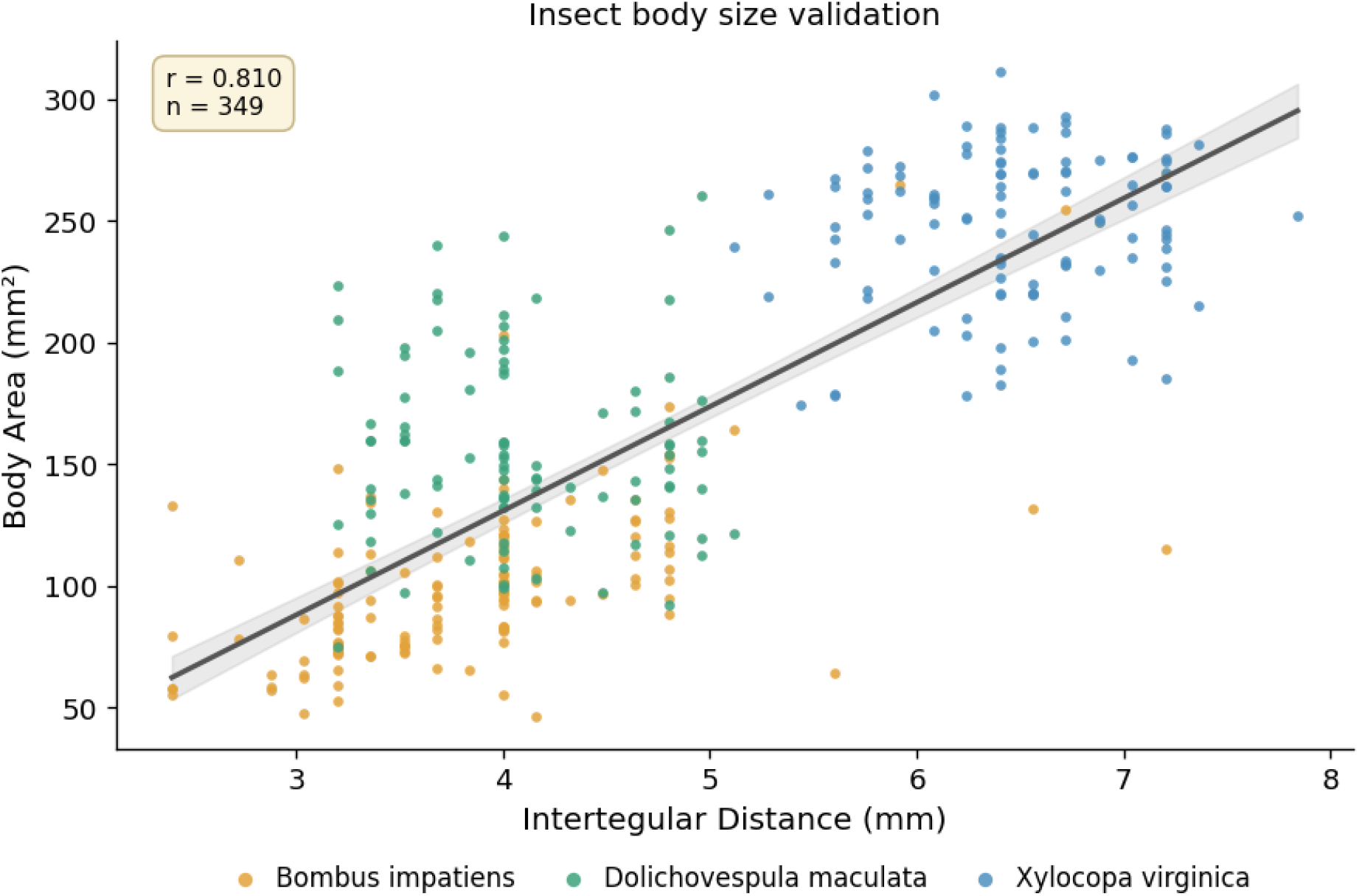
Relationship between hand-measured intertegular distance (ITD) and EcoMorph-derived body area across three bee species (*Bombus impatiens, Dolichovespula maculata, Xylocopa virginica*). Each point represents an individual specimen. The solid line indicates the pooled ordinary least squares regression with 95% confidence interval (shaded). r = Pearson correlation coefficient; n = total specimens.

EcoMorph counted insects almost perfectly from an insect box across a wide range of densities (7-80 specimens per image; mean 31, n = 12; Figure 9). Predicted counts exactly matched manual counts in 9 of 12 images, with the remaining three differing by a single insect (RMSE = 0.50, mean absolute error = 0.25 insects). The regression was near-identity (slope = 0.99, intercept = 0.39, R^2^ = 1.000), and residuals were symmetric around zero with no trend across the count range (Figure 9B), indicating no systematic count bias.

**Figure 9.**
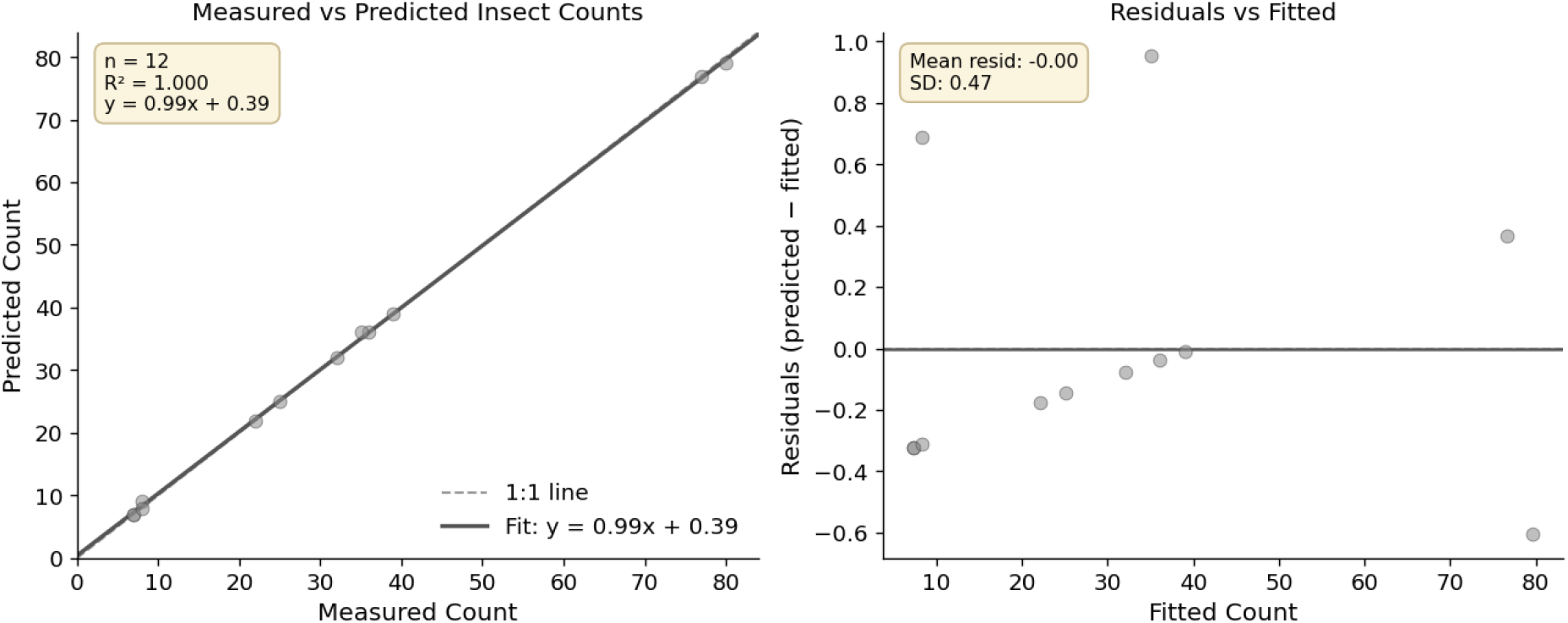
Validation of EcoMorph’s insect count function against manual counts (n = 12 images, 7-80 specimens per image). (A) Predicted versus manual counts. The solid line is the fitted regression (R^2^ = 1.000); the dashed line indicates the 1:1 relationship. (B) Residuals (predicted − manual) versus fitted counts, showing errors distributed symmetrically around zero (mean = 0.0, SD = 0.47) with no systematic trend across the count range. All residuals fell within ±1 insect.

## 4. Discussion

### 4.1 Summary of findings

EcoMorph demonstrated strong agreement with manual measurements across three measurement types (i.e., area, linear dimensions, and object count) spanning two ecological scales. For intermediate-scale floral area, the system agreed closely with manual ImageJ measurements under simple-background conditions (R^2^ = 0.935) and maintained strong accuracy under complex backgrounds with variable lighting, partial occlusion, and dense vegetation (R^2^ = 0.928), generating valid predictions for 95% of images. For fine-scale insect morphometrics, EcoMorph-derived body area was strongly correlated with hand-measured intertegular distance (r = 0.810, n = 349), capturing body-size variation consistent with a widely used morphometric standard across species spanning a broad size range (Bombus impatiens, Dolichovespula maculata, and Xylocopa virginica). For object counting, EcoMorph matched manual insect counts with almost perfect accuracy across a wide density range (7-80 per image; R^2^ = 1.000).

### 4.2 Comparison with existing morphometric tools

EcoMorph’s performance compares favorably with existing automated morphometric tools across both spatial scales and methodological approaches. At fine spatial scales, MAPHIS uses hierarchical image segmentation to extract detailed arthropod phenotypes but requires task-specific training for each application, limiting generalization across taxa (Mráz et al., 2024). At broader scales, classical computer vision workflows, such as those in Zhu et al. (2023), achieve high accuracy (R^2^ > 0.97) in measuring leaf morphology but rely on specialized lab imaging systems (e.g., the Scan1200), thereby limiting their field applicability. Deep learning (DL) approaches for plant phenotyping achieve>97% feature-identification accuracy under controlled greenhouse conditions (Pound et al., 2017), but performance typically degrades in field settings with variable lighting and complex backgrounds. Drone-based DL methods for floral surveys achieve strong agreement between automated and manual counts but show detection limitations in visually complex environments (precision 0.769, recall 0.849; Sookhan et al., 2024). EcoMorph’s ability to maintain R^2^ > 0.92 under complex-background conditions, while also generalizing to fine-scale insect morphometrics through the same prompt-based interface and consumer-grade imaging hardware, represents a key advance for ecological applications that require measurement across diverse taxa, imaging contexts, and accessibility constraints.

### 4.3 Limitations and future directions

A few limitations warrant consideration when applying EcoMorph to ecological research. First, SAM3 occasionally struggles with highly filamentous, small, or finely divided structures, as observed for certain flower morphologies in our validation. Researchers working with taxa exhibiting such structures should validate performance on representative samples before large-scale application.

Second, the computational requirements of SAM3 inference exceed those of lighter-weight DL models such as YOLO, potentially limiting throughput on very large image datasets. However, the improved accuracy, broader applicability, and elimination of task-specific training may justify additional computational costs for most research applications. Cloud-based processing and continued advances in model efficiency will likely reduce this constraint over time.

### 4.4 Conclusions

Morphological data on plant, animal, and habitat traits at different scales are used throughout basic and applied ecology. Collecting this trait data using traditional approaches is labor- and time-intensive. Computer vision and artificial intelligence offer opportunities for more consistent, standardized, and rapid data collection, but previous approaches have required the creation of large annotated image libraries to train these models. Using the Segment Anything Model 3 (SAM3) model and developing a modular pipeline with separate Object Segmentation, Morphology Extraction, and Measurement Scaling Modules, EcoMorph allows for flexible, customizable, and rapid analysis of a variety of morphological traits (area, linear dimension, and count data) across scales from a broad range of images, subjects, and contexts.

## Acknowledgements

The authors would like to thank Scott Diloreto for providing access to flowering plants in the Penn State Greenhouses and Sabrina Adler and Mehrdad Mahdavi for valuable discussions during the development of thistool. We thank Laura D. Porturas for supplying insect specimens from the Frost Entomological Museum at the Pennsylvania State University (GBIF ID: d044be1d-484b-4ecf-89b9-5b63054de960). The authors recognize the Penn State Institute for Computational and Data Sciences (RRID: SCR_025154) for providing access to computational research infrastructure within the Roar Core Facility (RRID: SCR_026424).

